# Navigating Pubertal Goldilocks: The Optimal Pace for Hierarchical Brain Organization

**DOI:** 10.1101/2023.08.30.555584

**Authors:** Hanna Szakács, Murat Can Mutlu, Giulio Balestrieri, Ferenc Gombos, Jochen Braun, Morten L. Kringelbach, Gustavo Deco, Ilona Kovács

**Affiliations:** Laboratory for Psychological Research, Pázmány Péter Catholic University, 1 Mikszáth square, Budapest 1088, Hungary; Semmelweis University Doctoral School, Division of Mental Health Sciences, 26 Üllői road Budapest 1085, Hungary; Institute of Biology, Otto-von-Guericke University, 44 Leipziger Straße; Center for Behavioral Brain Sciences, Otto-von-Guericke University, 44 Leipziger Straße 39120 Magdeburg, Germany; Center for Brain and Cognition, Computational Neuroscience Group, Universitat Pompeu Fabra, 25-27 Ramon Trias Fargas Barcelona 08005, Spain; HUN-REN-ELTE-PPKE Adolescent Development Research Group, 1 Mikszáth Kálmán square Budapest 1088, Hungary; Centre for Eudaimonia and Human Flourishing, Linacre College, University of Oxford, Oxford, UK; Department of Psychiatry, University of Oxford, Oxford, UK; Center for Music in the Brain, Department of Clinical Medicine, Aarhus University, Aarhus, Denmark; Department of Information and Communication Technologies, Universitat Pompeu Fabra, 122-140 Carrer de Tànger, Barcelona 08018, Spain; Institució Catalana de la Recerca i Estudis Avançats (ICREA), 23 Passeig de Lluís Companys Barcelona 08010, Spain; Institute of Psychology, Faculty of Education and Psychology, Eötvös Loránd University, 25-27 Kazinczy street, Budapest 1075, Hungary

**Keywords:** brain development, electrophysiology, thermodynamics, entropy production

## Abstract

Adolescence is a timed process with an onset, tempo, and duration. Nevertheless, the temporal dimension, especially the pace of maturation, remains an insufficiently studied aspect of developmental progression. This study focuses on the modifications due to the different timings of developmental shifts during adolescence and addresses the impact of adolescent maturation on brain development. To reveal potential relationships between pubertal pace and the advancement of brain organisation, we analyse the connection between skeletal age-based maturation stages and hierarchical organisation in the temporal dynamics of resting-state EEG recordings (alpha frequency range). By adopting skeletal maturity as a proxy for pubertal progress and employing entropy production to measure hierarchical brain organisation, our findings indicate that an average maturational trajectory optimally aligns with cerebral hierarchical order. Adaptive developmental plasticity may not fully compensate for accelerated or decelerated timelines, potentially increasing the risk of behavioural problems and psychiatric disorders consequent to such alterations.

## Main

Similar to early childhood development, adolescence is a time-bound event with an onset, tempo, and duration. While the interaction between genetic and environmental factors is established as a fundamental contributor to diversities in adult behavioural consequences, the temporal aspect, specifically, the speed of maturation, remains an insufficiently explored feature of developmental progression^1,2^. In the current study, we focus on the alterations of hierarchical brain organisation, related to different timings of pubertal maturation.

Adolescent cerebral maturation involves intricate sequences of neurodevelopment^3,4^, interwoven with alterations by age and stress^5–7^. Normative trajectories delineating volumetric^8^ and synaptic density^9^ shifts underscore the ongoing developmental dynamics of the adolescent brain. It is important to acknowledge, however, that the comprehensive volumetric transformations and mean synaptic density reflect the cumulative outcome of anatomical reconfiguration within the human brain, where the different components undergo heterochronous maturation^10,11^. Increasingly earlier onset of puberty referred to as worldwide secular trends^12–14^, or delayed onset due to, e.g., malnutrition or eating disorders^15,16^ might have significant consequences on developmental trajectories, impacting a wide range of physiological and psychological processes. For instance, puberty-associated reductions in cortical thickness^17^ or synaptic density^18^ may manifest prematurely or belatedly, deterring the adaptability of regions maturing at later stages to the atypical developmental milieu. Premature or postponed puberty can also generate challenges in behavioural adjustment, potentially leading to social isolation and heightened anxiety, particularly in the absence of adequate coping mechanisms^19^, thereby emphasising the critical role of adolescence in mental health^8,20^. Consequently, it is essential to investigate the impact of pubertal timing on the developing brain.

Uncertainty around the impact of atypical pubertal timelines arises from ineffective maturity measures and difficulty dissociating chronological age from pubertal maturity^21,17,22–24^. While puberty onset and its potential impact on developmental trajectories have long been a topic of interest, with significant individual variability suspected^23,25,26^, a precise assessment of the relationship between maturity and brain development remains noticeably absent from the literature, which is due to the lack of a reliable method to assess maturity levels. The Tanner Scale^27–29^, considered as the benchmark, relies on subjective evaluations of physical attributes such as breast and testicle size, rendering it susceptible to evaluator subjectivity. This scale originates from a post-war longitudinal study carried out within an orphanage during 1949-1971, thus failing to account for contemporary nutritional conditions and secular trends in growth^13,28,30–32^. Self- and parent-report versions of this method have also been deemed unreliable^29,28,33^.

Recently, we have introduced ultrasonic bone age as a promising alternative for maturity assessments in human developmental research^34^, with evidence for selective maturity-dependent effects in cognitive^34,35^, motor^36^, and emotional development^37^. Operating without harmful radiation, this technology gauges acoustic conductivity at hand and wrist growth zones (*Fig. 1a*) to estimate bone age, demonstrating robust correlation with pubertal hormone levels^28,33,38,39^, X-ray estimations^40^, and displaying high reliability^40^. This method overcomes traditional pubertal staging accuracy issues by providing an objective, continuous measure with unprecedented accuracy. In the current study, skeletal maturity serves as a proxy for individual pubertal maturity status. Addressing the impact of pubertal timing on brain development, the degree of maturity of each participant is calculated based on the difference between their bone age (BA) and chronological age (CA). It is widely acknowledged that the timing of puberty can differ significantly among individuals and follows a normal distribution^41–44^, therefore, to allow for statistical comparison across groups with different maturity levels, in our cross-sectional study, we selected an equal number of decelerated (BA < CA), average (BA = CA) and accelerated (BA > CA) maturity participants after screening them for bone age (*Fig. 1*, please also consult *Fig. 2a* under the *Results* section, and the *Methods/Participants* section for details). This approach not only disentangles chronological age from pubertal maturity (*Fig. 2b*) but also provides insights into how adolescents with varying maturity levels navigate pubertal goldilocks.

**Fig. 1:**
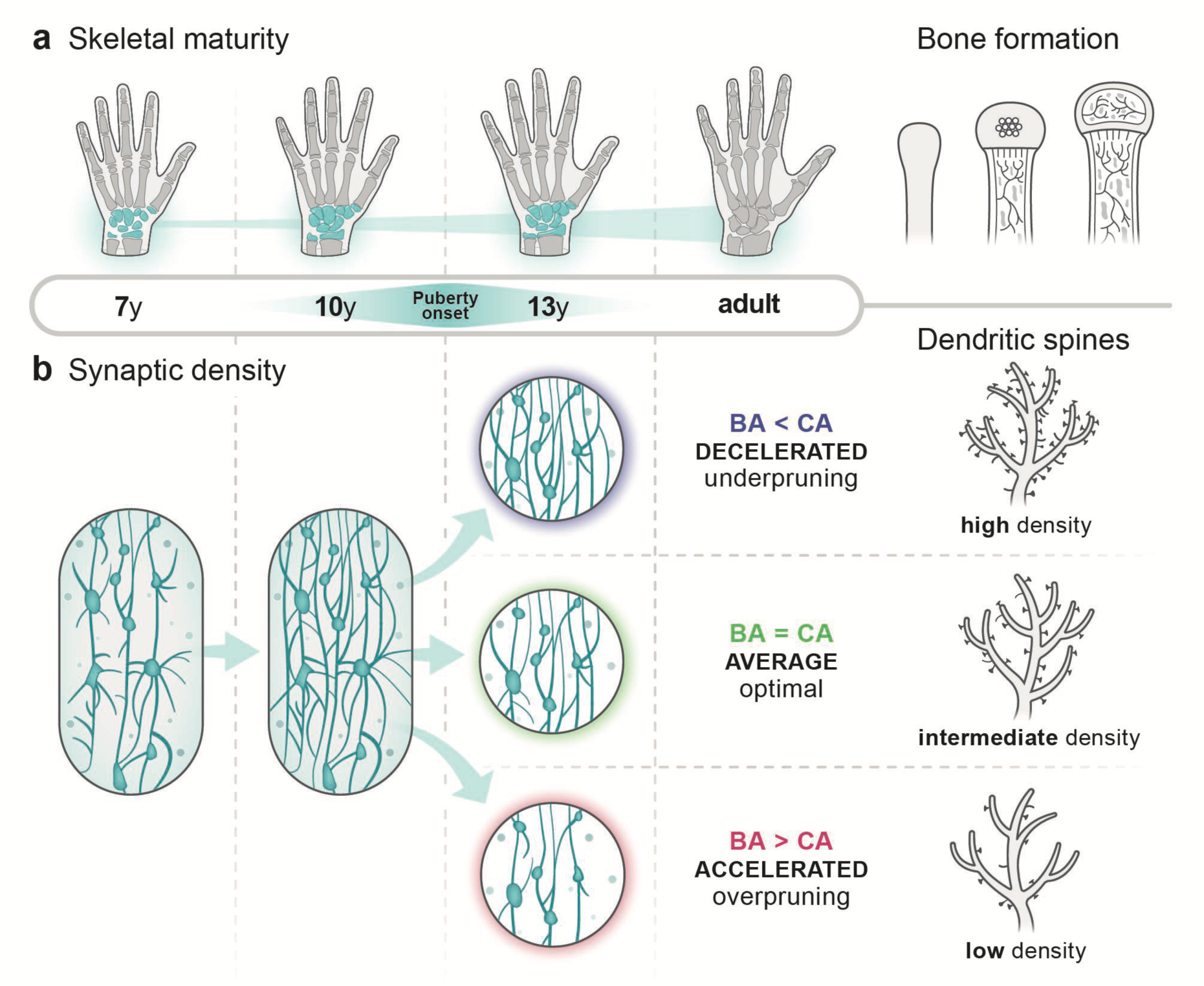
Anatomical aspects of skeletal and brain maturation. **a**, Bone development exhibits strong correlation with pubertal hormone levels. As bones form, acoustic conductivity within the growth zones undergoes alteration. This change in conductivity serves as the foundation for contemporary ultrasonic bone age assessments - a method offering objective, non-invasive estimation of biological age (BA) in children and adolescents. While a large proportion of the population aligns their BA with chronological age (CA), there are also accelerated individuals whose BA exceeds their CA, and decelerated ones, whose BA is less than their CA (the illustration does not show these variations, only examples of average bone development). **b**, Brain development is also profoundly influenced by the dynamic shifts in pubertal hormone levels. For example, a targeted process of synaptic pruning (reduction in dendritic spine density) is initiated by puberty onset, and pruning then continues until adulthood throughout the nervous system. However, as our conceptual figure suggests, synaptic pruning may not adhere to its expected course, but it may undergo dysregulation in cases of accelerated or decelerated pubertal maturation. Untimely maturation with dysregulated synaptic pruning may lead to the emergence of alterations in hierarchical brain organisation, which is the main issue addressed in the current study.

**Fig. 2.**
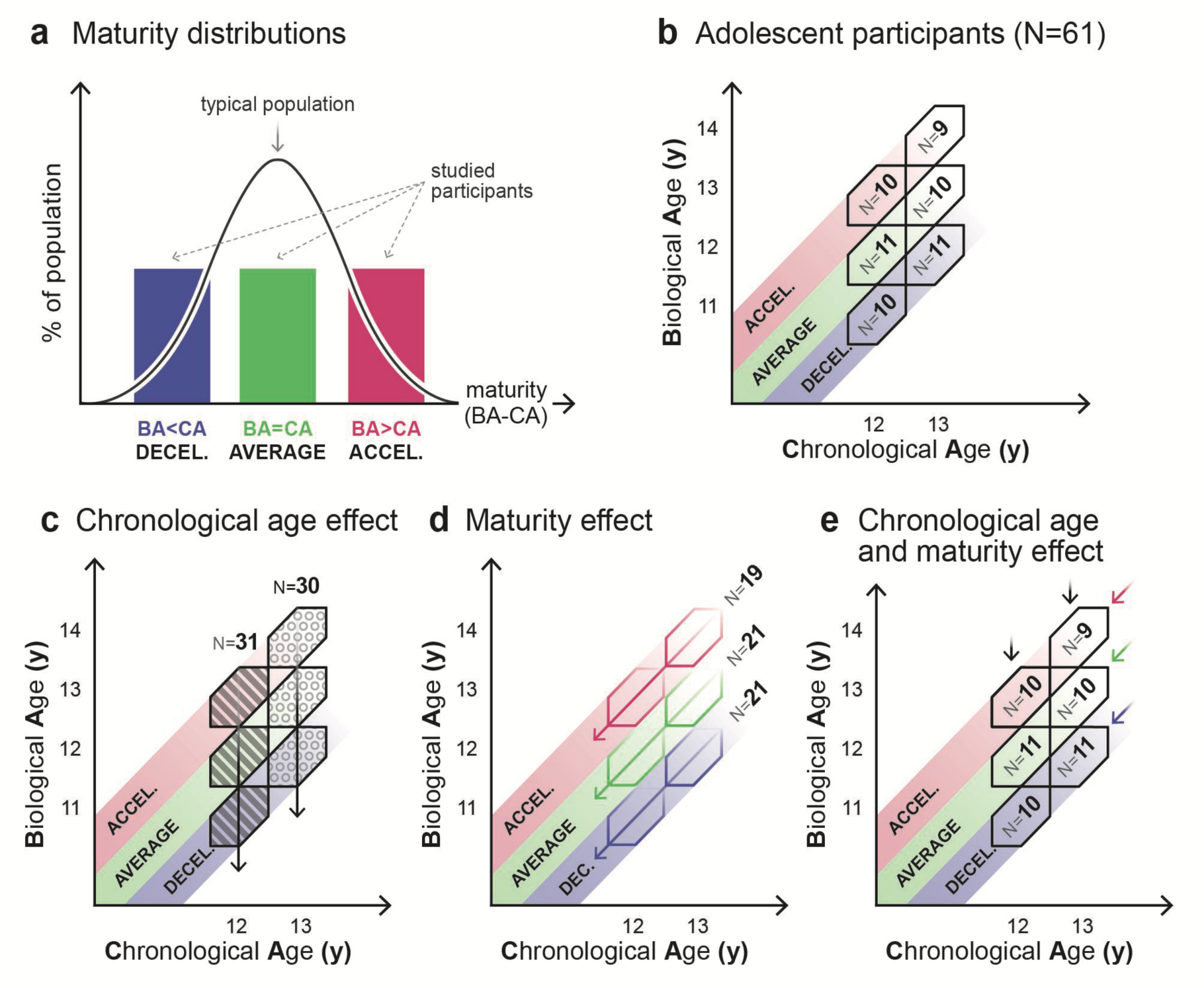
Participant grouping by adolescent age and maturity. **a,** The black curve illustrates the normal distribution of diverse maturity levels in a standard population, where maturity is represented along the horizontal axis as the discrepancy between biological age (BA) and chronological age (CA). Positive differences indicate accelerated, while negative differences indicate decelerated maturity. In the present study, to explore the influence of varying maturity levels, we initially assessed bone age using an ultrasonic device to estimate the biological age of participants. We then selected an approximately equal number of participants with decelerated, average, and accelerated maturity levels. **b,** Dissociation of biological and chronological age is achieved through the assignment of participants into one-year-wide hexagonal bins which ensures the absence of overlap between maturity groups. The data were then analysed through various grouping approaches: **c**, The effect of chronological age is analysed by averaging across different maturity groups within specific age groups; **d**, The effect of maturity is analysed by averaging across age groups within distinct maturity groups; **e**, By comparing data at the bin-level, the interplay between age and maturity effects is revealed.

As illustrated in *Fig. 1*, beyond bone development, the refinement of brain structure is also part of the anatomical changes associated with puberty and adolescence. One of the significant maturational events during this phase is the targeted elimination of synapses, known as synaptic pruning^45–48^ (*Fig. 1b*), which occurs in various species^49,50^ and across the nervous system^45^ during development to adapt to environmental conditions^48,50^. This process is vital for optimal brain function and shaping a healthy adult brain^47,51^. The timing of synaptic pruning has been linked to puberty onset in rodents^52,53^ and humans^18^, suggesting that early or late puberty may lead to a correspondingly early or delayed initiation of synaptic pruning. However, it is not clear whether synaptic pruning follows a normal course or becomes dysregulated when it begins earlier or later (*Fig. 1b*). As insufficient synaptic pruning or „underpruning” has been tied to neurodevelopmental disorders^4,22^ and excessive pruning or „overpruning” to psychiatric conditions such as schizophrenia^54,55^, it is important to investigate whether untimely maturation may lead to the emergence of alterations in brain organisation, thereby contributing to the current psychological challenges faced by adolescents^56–60^.

The temporal aspect of synaptic pruning holds significance within hierarchical neurodevelopment as well. Typically, transmodal association regions undergo peak synaptic density later than sensory zones, experiencing pruning throughout adolescence^11,45,61^. This design permits a broad experience-dependent refinement window but also introduces the prospect of prolonged vulnerability^11,62^. It is also becoming increasingly clear that developmental alterations in the functional^63,64^ and evolutionary^65^ hierarchies of the brain appear to mirror anatomical maturation patterns. Consequently, our primary objective is to investigate the precise influence of pubertal maturation on the configuration of associative brain regions, which manifest the most substantial developmental shifts during adolescence. For this purpose, we assess resting-state brain activity and extract the degree of hierarchical arrangement among participant groups exhibiting diverse levels of physical maturation.

Recent progress in neuroimaging has enabled the creation of normative brain-growth charts^8,66^, similar to those applied for anthropometric attributes like height and weight. Despite their reliance solely on chronological age and their predominant cross-sectional nature, these investigations reveal the spatiotemporal dynamics of developmental plasticity and have even identified a sensorimotor-to-associative sequence of refinement^10,11^. The latter findings align with the progress of phylogenetic brain development from less variable unimodal areas towards heteromodal association cortices^65,67,68^. It is important to recognise that the most recently evolved associative networks within the brain might be particularly vulnerable to diverse pubertal timelines, given their ontogenetic maturation coincides more closely with the adolescent phase. Irregular timelines could also relate more to alterations in the finely tuned intrinsic activity of these networks, as opposed to extensive anatomical reconfigurations. While fMRI has proven useful in assessing intrinsic resting-state activity, it is sensitive to neural, vascular, and respiratory influences. Therefore, direct measures of neural activity might offer better options for investigating this intrinsic behaviour. Resting-state EEG provides direct measures of intrinsic cortical activity, allowing for the detection of changes in oscillatory activity and functional connectivity that are not captured by fMRI. In this study, we rely on resting-state EEG in the alpha frequency range (see *Methods*) to compare the progression of hierarchical brain organisation across groups with early, on-time and late maturing groups as defined by the difference between their bone age and chronological age.

To reveal any relationships between pubertal pace and the advancement of brain organisation, we analyse how skeletal age-based maturity levels link to high-level features of resting-state EEG-dynamics. Specifically, we introduce “entropy production” to capture the degree of functional hierarchical organisation of the brain. In thermodynamics and systems biology, the asymmetry and directionality of flow in the state space of the components of a living system is known as “breaking of the detailed balance” and give rise to non-equilibrium states which can be captured by the level of non-reversibility in time^69,70^. In other words, different levels of hierarchical organisation allow the orchestration of the whole-brain dynamics accordingly in different ways. Hierarchy is built up by the underlying asymmetry in the directionality of information flow between different regions. Thermodynamics establishes a direct link between hierarchy, non-equilibrium, and time irreversibility. This comes from the fundamental idea of the second law of thermodynamics, explicitly stating that a system will go from order to disorder over time. We capture these different levels of organisation and non-equilibrium through the level of irreversibility, i.e., through the “arrow of time.”^69,71^

Asymmetric flow can be observed in state spaces at various scales, from the molecular^72^ to the macroscopic brain dynamics^70^. In general, asymmetric flow reflects a degree of non-reversibility in the dynamics of the system. In the case of the brain, such non-reversibility is thought to reflect the hierarchical organisation of brain activity^69,73^. Entropy production is a direct measure of non-reversibility and indirectly captures the functional hierarchy expressed by brain activity^69,71,74^ (see *Methods*). The purpose of our study is to investigate the relationship between entropy production, as a proxy of hierarchical brain activity, and the level of physical maturation.

## Results

As it is explained in *Fig. 2b*, we dissociated biological and chronological age through the assignment of participants into one-year-wide hexagonal bins to ensure the absence of overlap between maturity groups with approximately equal numbers of participants. We then analysed the data through various grouping approaches illustrated in *Fig. 2c-d*, allowing for the analysis of chronological age and maturity levels independently. Results are discussed according to those groupings. We have obtained and analysed closed-eye resting state high-density EEG recordings in the alpha frequency range (8-12 Hz) in 61 adolescent, and 26 emerging adult female participants (see *Fig. 2* and *Methods/Participants*). To measure the level of hierarchical brain activity in resting-state EEG recordings, we parsed the recorded neural data into 12 topographically distinct, phase-based activity patterns, referred to as “brain states” (see *Fig. 6*), utilising a modified *k*-means clustering algorithm^75^. Each brain state captures patterns where two major electrode sets record congruent signals, but these sets are in opposition to one another. After extracting the brain state patterns, we studied the sequence in which these brain states occurred, specifically focusing on the asymmetrical nature of their transitions. A sequence of consecutive brain states is illustrated in *Fig. 7a* as a representative example in the *Methods*/*Data Analysis* section, while *Fig. 7b* presents a table that counts the observed occurrences of each transition within this example, highlighting their asymmetric quality.

### Chronological age related entropy production

To evaluate the effect of chronological age on entropy production, we averaged across maturity groups within a 12-year-old (N=31, mean age=12.48y, SD=0.27y) and a 13-year-old (N=30, mean age=13.58y, SD=0.29y) group. In order to have normative data from young adults, we also included 22-year-old participants (N=26, mean age=21.69y, SD=0.64y). For further details see *Fig. 2c*, *Fig. 3c*, and *Methods*/*Participants*. Importantly, adult data should only be compared to age-averaged results of adolescents as the adult cohort has not been selected with respect to maturity. Given that bone formation is completed by adulthood, bone age assessment is irrelevant in adults. Therefore, adult data were included in the statistical analysis only where the basis of the analysis is chronological age, irrespective of maturity.

**Fig. 3.**
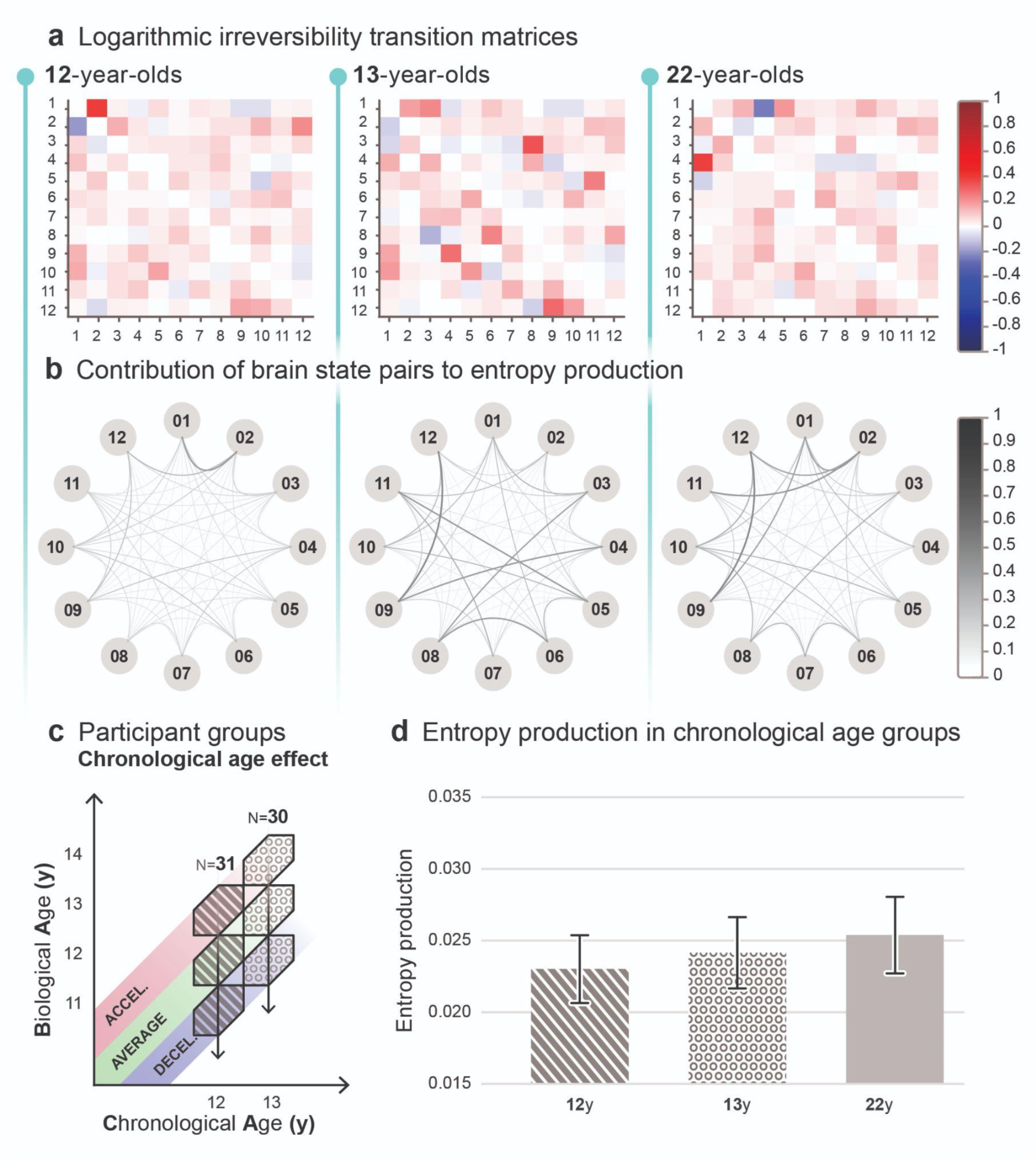
Chronological age related entropy production. **a**, Logarithmic irreversibility transition matrices illustrate two distinct, non-symmetric values in each brain state pair, derived from the log ratio of the forward and reverse transition probabilities. The matrices inform about the preference for directionality in brain state switching. The extent of deviation from 0 reflects the strength of directional preference (forward transitions in red, backward transitions in blue), indicating the level of irreversibility and hierarchical organisation in the brain. The relative similarity in the variability in hue and intensity indicates comparable entropy production between age groups. **b**, Graphs represent the cumulative contribution to entropy production of both forward and reverse transitions and reveal the level of irreversibility in the dynamics of each brain state pair within age groups. Nodes represent the 12 brain states, edges indicate pairwise contributions to entropy production, where increasing thickness and darkness represent greater contribution, therefore larger irreversibility. There seems to be a subtle rise in contributions to entropy production with age. **c**, The effect of chronological age is analysed by averaging across different maturity groups within specific age groups. **d**, Bar plots illustrate the average entropy production and respective 95% confidence interval for each age group calculated via inverse-variance weighting. Differences among groups were statistically assessed using two-sample z-tests, accompanied by Hedges’ g effect size calculations, yielding no statistically significant difference among the age groups in pairwise comparisons (*p*>0.05) and small effect sizes in all comparisons.

*Fig. 3a* and *3b* provide a detailed look at brain state dynamics within each age-group. Logarithmic irreversibility transition matrices in *Fig. 3a* illustrate a contribution to entropy production of brain state transitions in age groups, derived from the log ratio of the forward and reverse transition probabilities, estimated from transition counts. Matrices were generated through averaging individual matrices within each age group, followed by normalisation of values to a -1 to +1 range (using the largest absolute value of the global extremum), indicated by the colour scale. Each matrix holds 12×12 values, with the starting brain states on the horizontal and the subsequent states in brain state switching on the vertical axis. Log ratios of the forward and reverse transition probabilities are non-symmetric. Positive and negative values indicate a preference for forward and reverse transitions, respectively. The relative similarity in the variability in hue and intensity indicates comparable entropy production between age groups, with a slight indication of age-related improvement. Graphs in *Fig. 3b* show total entropy production contributions of brain state pairs, computed by summing the log ratio values of forward and reverse transition probabilities within each pair, creating symmetric matrices. Graph values are generated by averaging the individual symmetric matrices within age groups, then normalising them between 0 and 1. The nodes represent the 12 brain states, with the most dominant state (highest percentage of total activity) positioned at the 12 o’clock position, and subsequent states are arranged clockwise in order of descending dominance. The edges indicate pairwise contributions to entropy production, where increasing thickness and darkness represent greater contribution. The slight increase in line thickness and darkness across age-groups may indicate a subtle rise in contributions to entropy production with age.

Group averages and variances of entropy production were computed via inverse-variance weighting to minimise the biassing effect on the weighted arithmetic mean introduced by variance. 95% confidence intervals of group-level entropy production were also calculated using the weighted group average. Weighted group averages revealed a tendency of increase in entropy production with age, as it is demonstrated in *Fig. 3d*. Specifically, the 12-year-old group exhibited an average mean entropy production of 0.02300 bit (95% CI: 0.02064-0.02537), while the average mean entropy production of the 13-year-old group was 0.02415 bit (95% CI: 0.02166-0.02663), and the average mean entropy production of the 22-year-old group was 0.02538 bit (95% CI: 0.02271-0.02805). To determine whether the group differences are statistically significant, we utilised two-sample z-tests to compare the group averages (see *Methods* for further details). While a slight increase in groupwise average entropy production with chronological age is observed, pairwise comparison of group averages were not statistically significant (*p*>0.05 in each comparison). Hedges’ g effect size estimates revealed small effect sizes in all cases: between the 12-year-old and 13-year-old (*g*=0.17), and between 13-year-old and 22-year-old (*g*=0.18) groups, and in the comparison of the 12-year-old and 22-year-old group (*g*=0.35).

Our statistical results indicate that, while there is a slight increase in entropy production with increasing chronological age, this variable independently does not have a substantial impact on entropy production, further warranting the inspection of maturity in adolescence.

### Maturity related entropy production

To analyse the relationship between maturity and group-level average entropy production, we sorted adolescent participants according to their level of maturity, irrespective of chronological age, forming three distinct groups: decelerated maturity (CA-1.5≤BA<CA-0.5; N=21, mean age=12.98y, SD=0.60y, mean BA-CA=-0.92, SD=0.24), average maturity (CA-0.5≤BA≤CA+0.5; N=21, mean age=12.97y, SD=0.60y, mean BA-CA=0.03, SD=0.29), and accelerated maturity (CA+1.5≥BA>CA+0.5; N=19, mean age=13.12y, SD=0.68y, mean BA-CA=0.88, SD=0.31). Participant groups are illustrated both in *Fig. 2d* and *Fig. 4c*.

**Fig. 4.**
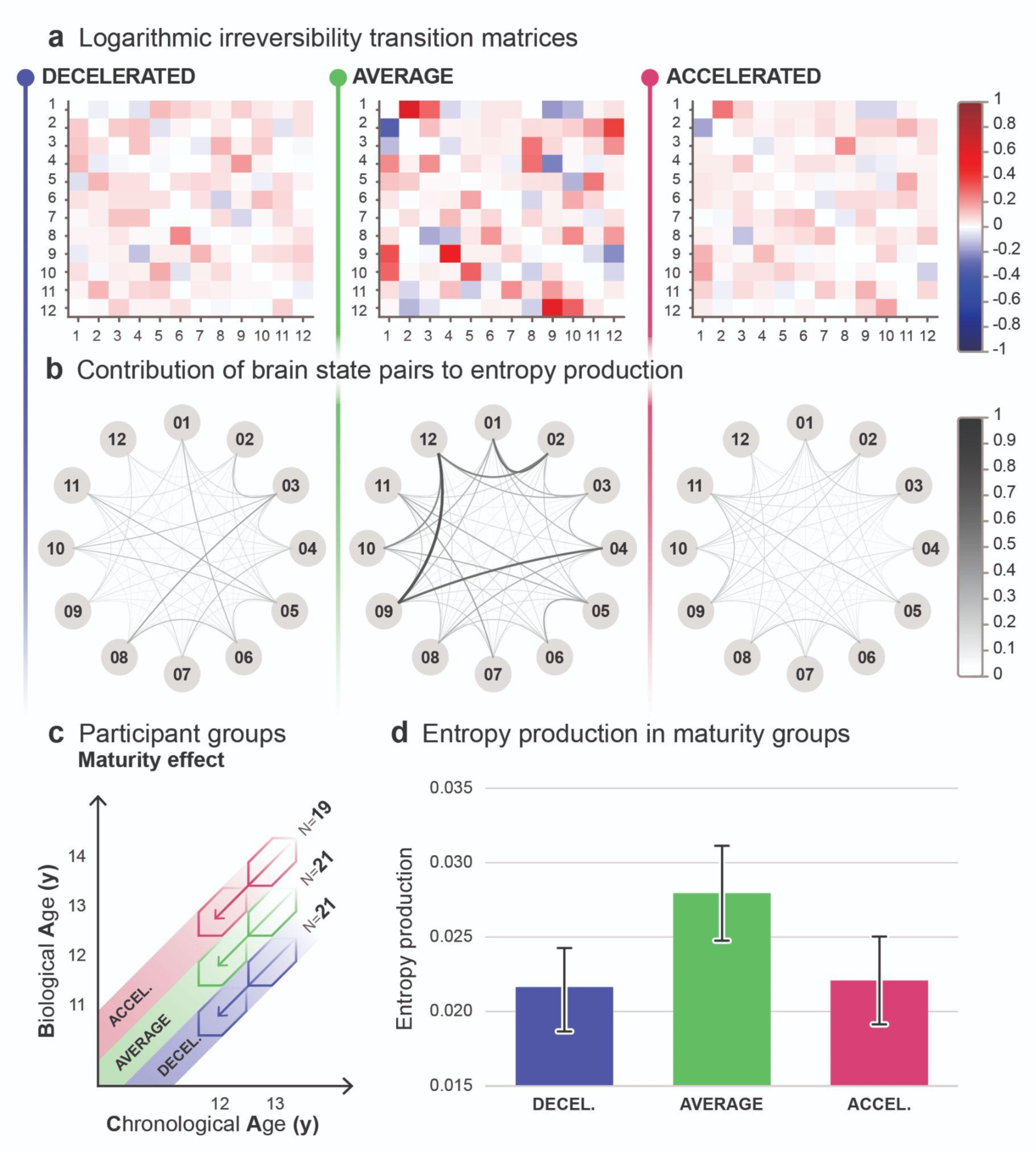
Maturity related entropy production. **a**, Logarithmic irreversibility transition matrices illustrate two distinct, non-symmetric values in each brain state pair, derived from the log ratio of the forward and reverse transition probabilities. The matrices inform about the preference for directionality in brain state switching. The extent of deviation from 0 reflects the strength of directional preference (forward transitions in red, backward transitions in blue), indicating the level of irreversibility and hierarchical organisation in the brain. The larger hue and intensity variation in the matrix of the average maturity group indicates higher entropy production relative to the decelerated and accelerated groups. **b**, Graphs represent the cumulative contribution to entropy production of both forward and reverse transitions and reveal the level of irreversibility in the dynamics of each brain state pair within maturity groups. Nodes represent the 12 brain states, edges indicate pairwise contributions to entropy production, where increasing thickness and darkness represent greater contribution, therefore larger irreversibility. Graphs indicate higher entropy production in the average maturity group, apparent in the thicker and darker lines. **c**, The effect of maturity is analysed by averaging across age groups within distinct maturity groups. **d**, Bar plots illustrate the average entropy production and respective 95% confidence interval for each maturity group, calculated via inverse-variance weighting. Using two-sample z-tests, we compared the average group to both decelerated (*p*=0.0028) and accelerated (*p=*0.0082) groups, observing significant differences in each comparison with large effect sizes (Hedges’ g).

*Fig. 4a* and *4b* offer detailed insight into brain state dynamics within each maturity group. As it is described above, with respect to *Fig. 3a*, logarithmic irreversibility transition matrices in *Fig. 4a* illustrate a contribution to entropy production of brain state transitions, in this case within decelerated, average, and accelerated maturity groups among adolescent participants. The larger hue and intensity variation in the matrix of the average maturity group indicates higher entropy production relative to the decelerated and accelerated groups. Graphs in *Fig. 4b* represent the total contribution of both forward and reverse transitions to entropy production (see the description related to *Fig. 3b* above), indicating higher entropy production in the average maturity group, apparent in the thicker and darker edges of the graph compared to that of the decelerated and accelerated groups.

Using inverse-variance weighting, we calculated the group-level average and variance of entropy production to minimise the distorting effect of variance, and determined the 95% confidence intervals based on the weighted group average. The inverse-variance weighted group average and confidence interval of the decelerated maturity (mean=0.02146 bit, 95% CI: 0.01866-0.02426), average maturity (mean=0.02794 bit, 95% CI: 0.02475-0.03113), and accelerated maturity (mean=0.02208 bit, 95% CI: 0.01913-0.02503) groups are illustrated in *Fig. 4d*. It is clear that the average maturity group expresses higher average entropy production than both accelerated and decelerated maturity groups. We utilised two-sample z-tests to determine the statistical significance of the apparent differences, reported with Hedges’ g effect size tests. Results showed that the average maturity group expresses significantly higher groupwise average entropy production than both accelerated (*p*=0.0082, *g*=0.83) and decelerated (*p*=0.0028, *g*=0.92) maturity groups. Effect sizes were large in both pairwise tests.

The average maturity group consistently expressed significantly higher (*p*<0.05) average entropy production than both decelerated and accelerated groups with additional clustering approaches, specifically, working with 6, 8 and 10 brain states, and 12 brain states that are specific to each unique group; see the Open Science Framework link under *Data availability* for further details.

Our findings show that maturity significantly influences entropy production, irrespective of chronological age, with the group-level entropy production being significantly higher in the average maturity group.

### Chronological age and maturity related entropy production

To study the interplaying effect of chronological age and maturity, we divided adolescents into two age brackets, namely 12-year-olds (N=31, mean age=12.48y, SD=0.27y) and 13-year-olds (N=30, mean age=13.58y, SD=0.29y), and within these brackets, we further assorted participants into three distinct maturity groups, forming six groups, referred to as bins, in total.

In the 12-year-old group, we identified three maturity subgroups: decelerated maturity (CA-1.5≤BA<CA-0.5; N=10, mean age=12.41y, SD=0.26y, mean BA-CA=-0.86, SD=0.21); average maturity (CA-0.5≤BA≤CA+0.5; N=11, mean age=12.48y, SD=0.28y, mean BA-CA=0.04, SD=0.31); and accelerated maturity (CA+1.5≥BA>CA+0.5; N=10, mean age=12.54y, SD=0.27y, mean BA-CA=0.91, SD=0.34). Similarly, in the 13-year-old group, we established three maturity subgroups: decelerated maturity (CA-1.5≤BA<CA-0.5; N=11, mean age=13.49y, SD=0.24y, mean BA-CA=-0.97, SD=0.27); average maturity (CA-0.5≤BA≤CA+0.5; N=10, mean age=13.52y, SD=0.31y, mean BA-CA=0.03, SD=0.28); and accelerated maturity (CA+1.5≥BA>CA+0.5; N=9, mean age=13.76y, SD=0.26y, mean BA-CA=0.85, SD=0.28). Participant bins are illustrated both in *Fig. 2e* and in *Fig. 5c*.

**Fig. 5.**
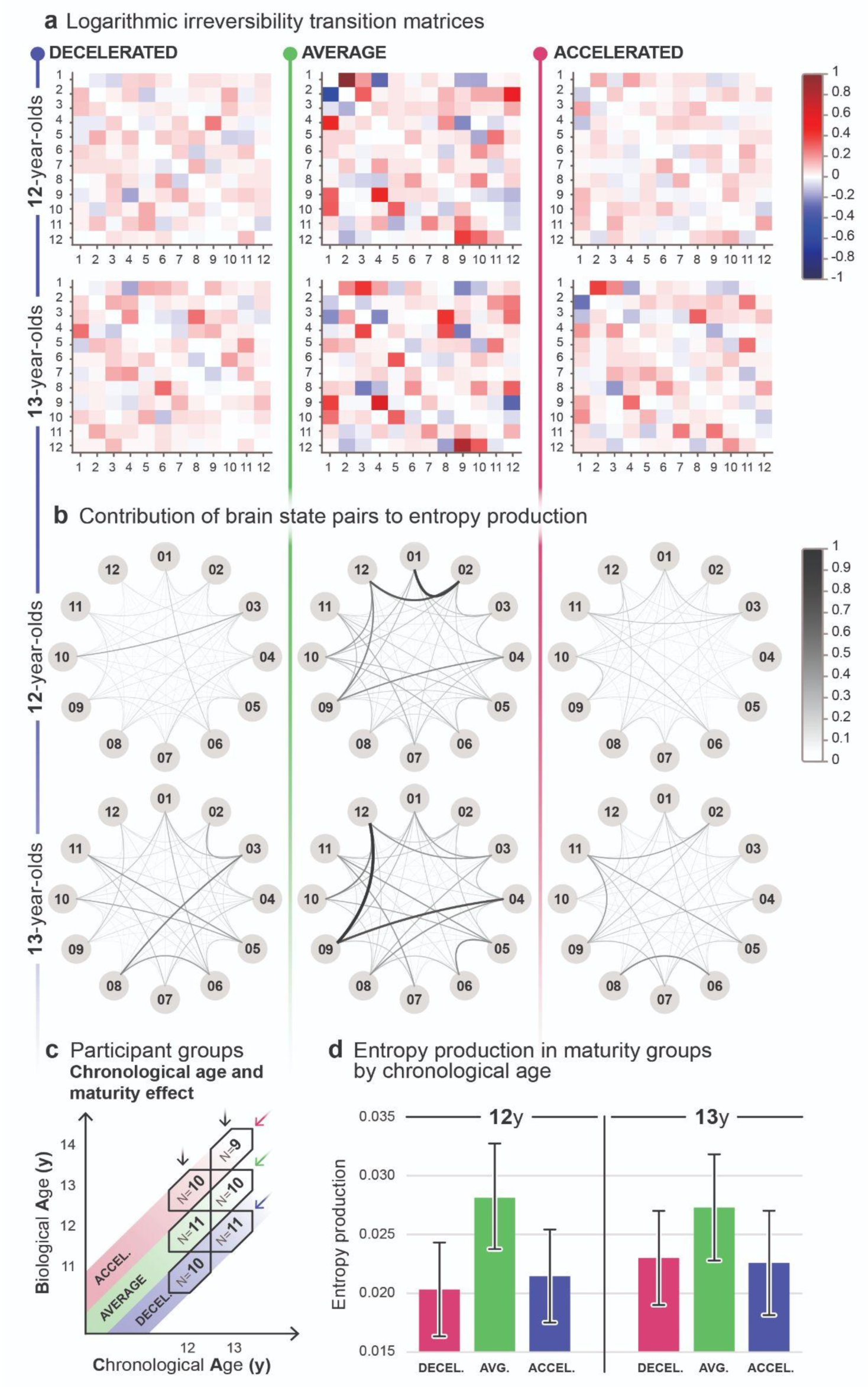
Chronological age and maturity related entropy production. **a**, Logarithmic irreversibility transition matrices illustrate two distinct, non-symmetric values in each brain state pair, derived from the log ratio of the forward and reverse transition probabilities. The matrices inform about the preference for directionality in brain state switching. The extent of deviation from 0 reflects the strength of directional preference (forward transitions in red, backward transitions in blue), indicating a higher level of irreversibility and hierarchical organisation in the brain. The hue and intensity diversity in the matrices in the average maturity groups across both age groups indicates higher entropy production values compared to decelerated and accelerated maturity groups. **b**, Graphs represent the cumulative contribution to entropy production of both forward and reverse transitions and reveal the level of irreversibility in the dynamics of each brain state pair within age groups. Nodes represent the 12 brain states, edges indicate pairwise contributions to entropy production, where increasing thickness and darkness represent greater contribution, therefore larger irreversibility. Graphs indicate higher entropy production in both average maturity groups, apparent in the thicker and darker lines. **c**, The effect of maturity is analysed by averaging across age groups within distinct maturity groups. **d**, Bar plots illustrate the average entropy production and respective 95% confidence interval for each maturity group, calculated via inverse-variance weighting. Using two-sample z-tests, we compared the average group to both decelerated (*p*=0.0098) and accelerated (*p=*0.0261) groups, observing significant differences in each comparison with large effect sizes (Hedges’ g) in 12-year-olds, however, the analysis of 13-year-olds did not yield significant results.

*Fig. 5a* and *5b* offer detailed insight into brain state dynamics within each bin. As it is described above, with respect to *Fig. 3a*, logarithmic irreversibility transition matrices in *Fig. 5a* illustrate a contribution to entropy production of brain state transitions, in this case within decelerated, average, and accelerated maturity groups within age groups. In both age brackets, the larger hue and intensity variation in the matrix of the average maturity group indicates higher entropy production relative to the decelerated and accelerated groups. Graphs in *Fig. 5b* represent the total contribution of both forward and reverse transitions to entropy production (see the description related to *Fig. 3b* above), indicating higher entropy production in the average maturity group in both age groups, apparent in the thicker and darker edges of the graph compared to that of the decelerated and accelerated groups.

We utilised inverse-variance weighting to calculate group averages and variances of entropy production, minimising the biassing effect that variance introduces to the weighted arithmetic mean. Additionally, we determined the 95% confidence intervals for group-level entropy production based on the weighted group average. As it is suggested in *Fig. 5d*, the average maturity group expresses higher average entropy production at the group-level than decelerated and accelerated groups in the same chronological age category. We performed two-sample z-tests to determine the statistical significance of differences. Two-sample z-tests, accompanied by Hedges’ g effect size estimations, confirmed statistically significant differences among the 12-year-old maturity bins (see *Fig. 5d*); we compared the average entropy production of the average maturity (mean=0.02823 bit, 95% CI: 0.02374-0.03271) bin to both decelerated (mean=0.02032 bit, 95% CI: 0.01633-0.02431; *p*=0.0098, *g*=1.120) and accelerated (mean=0.02145 bit, 95% CI: 0.01750-0.02540; *p*=0.0261, *g*=0.963) bins, revealing significant differences and large effect sizes in these comparisons. Performing two-sample z-tests on the maturity bins of the 13-year-olds did not yield a significant result (*p*>0.05 in all comparisons).

Effect size measurements revealed a medium effect size (*g*=0.61) between the average maturity (mean entropy production=0.02728 bit, 95% CI: 0.02277-0.03179) and decelerated (mean entropy production=0.02299 bit, 95% CI: 0.01899-0.02700) bins, and a large effect size in the comparison of average versus accelerated (mean entropy production=0.02257 bit, 95% CI: 0.01813-0.02701; *g*=0.67) bins.

Our results show that maturity strongly influences entropy production among 12-year-old participants. While we did not find significant differences between the maturity groups in the 13-year-old age bracket, the higher group-level entropy production in the average maturity group is still present in the logarithmic irreversibility transition matrices (*Fig. 5a*), in the graphs of *Fig. 5b*, and in the effect size calculations, indicating a persistent pattern of increased entropy production associated with average maturity.

## Discussion

A novel question was addressed regarding the role of adolescent maturation in brain development. While individual differences in maturity among teenagers are well-established, it has been unclear whether there is a difference in brain development between groups of adolescents with diverse maturity levels. The introduction of skeletal age into research now allows for conclusive answers to this question in a field obscured by methodology issues around maturity assessments. Our results suggest that an average maturational trajectory might be optimal in terms of hierarchical brain organisation. The significance of this study stems from the possibility that developmental plasticity may not fully compensate for accelerated or decelerated timelines, and a heightened risk of behavioural problems and psychiatric disorders may arise from such alterations.

To address how well adolescents with different maturational pace can navigate pubertal goldilocks, we relied on skeletal age as a proxy for pubertal progress and resting-state EEG-derived entropy production as a gauge of hierarchical brain organisation. As illustrated in *Fig. 2*, we employed the strategy of selecting an equal number of participants with accelerated, average, and decelerated maturity levels. We have also completed our sample by a group of emerging adult participants who were not selected by maturity levels since bone age assessments are not informative at this age. The dissociation between biological age and chronological age illustrated in *Fig. 2b* allowed us to independently analyse the impact of age and maturity on the extent of resting-state EEG-derived entropy production.

First, we analysed the relationship between chronological age and entropy production by averaging across the three maturity groups within the 12 and 13-year-old participant groups (*Fig. 2c* and *Fig. 3c*) and comparing them to the emerging adult group. Despite a noticeable increasing trend in entropy production with age (*Fig. 3d*), chronological age in itself did not have a statistically significant effect on entropy production. The similar logarithmic irreversibility transition matrices of the age-groups (*Fig. 3a*), and the relatively similar dispersion of entropy production among brain state pairs (*Fig. 3b*) support the statistical results. These findings suggest that the level of hierarchy in the adolescent brain may not be meaningfully influenced by chronological age alone. It is important to note that we lack information about the adolescent maturational trajectories of emerging adult participants, while adolescents have been screened and selected for the maturity groups in an approximately equal number. Therefore, the distributions of individuals with different maturity levels might be different between the emerging adults and adolescents (*Fig. 2a*), and their direct comparison should be approached with caution.

To investigate the influence of maturity on entropy production, we categorised adolescent participants into three distinct groups based on their level of maturity (decelerated, average, and accelerated maturity groups in *Fig. 2d* and *Fig. 4c*.). Statistical analysis of group averages (*Fig. 4d*) revealed significantly higher entropy production in the average maturity group with large effect sizes, pointing at a considerably higher level of hierarchical organisation when chronological age and bone age align. A closer look at the logarithmic irreversibility transition matrices (*Fig. 4a*) illustrating the strength of directional preference in switching among the 12 states provides further insight into the role of maturity in hierarchical organisation. The average maturity group has a relatively higher number of brain state pairs among which a strong directionality is established. The contribution of brain state pairs also reflects an elevated level of hierarchy in the average maturity group (*Fig. 4b*). The dynamics between brain states underlying the entropy production levels in each group indicate well-established causal interactions, and hierarchical brain organisation in the average maturity group, while these relationships seem relatively less dense in the decelerated and accelerated groups. These findings indicate that physical maturity is inherently linked to the shaping of irreversible, causal interactions within the cortex^69^ during adolescence. Furthermore, it suggests that the optimal maturational pace for fostering these relationships is one in which chronological and biological age are closely matched.

To see whether the maturity related effect is stable across different chronological ages, we compared the three maturity groups within the 12- and 13-year-old age groups (*Fig. 2d* and *Fig. 5c*). The analysis revealed that the average maturity group displays significantly higher entropy production compared to both the decelerated and accelerated maturity groups among the 12-year-olds (*Fig. 5d*). While this tendency is still apparent in the 13-year-old group, statistically significant differences between maturity groups were not detected. However, the logarithmic irreversibility transition matrices of each bin (*Fig. 5a*) make a case for the optimal pace of maturation even in the absence of statistical significance, as the average maturity groups in both age brackets convey a larger number of brain state pairs where a strong preference of switching direction is established. The contribution of each brain state pair to entropy production (*Fig. 5b*) also highlights a larger number of well-established irreversible interactions in the average maturity groups. Taken together, the statistical results (Fig. 5d) may suggest that the clear advantage of an on-time maturational pace is limited to a specific chronological age window with a declining tendency in older age groups, however, this possibility may need further investigation as the graphical analyses presented in Fig. 5a and 5b suggest a persistent pattern of increased entropy production associated with average maturity.

To conclude, we would like to re-emphasise that maturation plays a pivotal role in shaping both brain structure and behaviour. This process ensures that the brain efficiently coordinates distributed computations across its entirety, allowing not only survival but also optimal functioning. This efficiency hinges on a hierarchical orchestration of state-dependent, self-organised brain activity over time. This orchestration facilitates near-optimal information processing and transfer while minimising energy consumption. Therefore, a paramount concern of a comprehensive theory of brain function is the exploration of hierarchy. This entails unravelling the dynamic and evolving self-organisation of brain states, spanning from stability to transitions, as they navigate through probabilistic and sometimes chaotic shifts. Here, we assumed that the inherent hierarchy of spontaneous brain activity might be reflective of the level of maturation. Harnessing the principles of thermodynamics, we quantified hierarchy by computing the level of irreversibility in brain signals during resting state conditions. Specifically, we captured the level of irreversibility through entropy production^69,71,74,76,77^ as computed through a sophisticated clustering strategy. Note that resting states cover a rich range of the dynamical repertoire of the brain and thus those are ideal to characterise the most general and wide variety of interaction across the whole brain, i.e., the underlying hierarchical orchestration.

In terms of limitations, the current project is confined to the examination of adolescent females, a limitation that might be apparent. However, a selection was essential when aiming to distinctly disentangle the impacts of maturation and chronological age on brain development. Involving both sexes would have made it difficult to carry out the detailed analysis within the narrow temporal windows of biological and chronological age depicted in *Fig. 2b*. As male pubertal onset occurs approximately 1-1.5 years later than that of females^32,78^, this analysis would necessitate a considerably larger sample size. We opted for inclusion of females since menarche age provides valuable supplementary data affirming the accuracy of bone age assessments (see *Methods*). Given the conclusive findings with females, underscoring a substantial role of biological age in brain development, it is plausible to assume a comparable effect might manifest in males. Indeed, we would expect that maturity is a relevant factor in the general brain development of males as well; however, owing to the distinct gonadal hormone levels governing female and male developmental trajectories^21,79^, disparities are also anticipated, including the timing of the impact of maturity levels.

An additional limitation of our study pertains to its cross-sectional design within a relatively narrow age spectrum, which prevents an exploration of potential long-term consequences due to variations in pubertal pace. The evolutionarily expanded heteromodal association cortices, mentioned in the introduction, hosting a range of higher order functions^11,65,67,68,80^ may have a potential to compensate when confronted with divergent maturational paces. Nevertheless, the consequences of accelerated or delayed development on relevant and precisely timed adolescent brain alterations such as cortical volume reduction^81,82^, grey matter thinning^17,83^, and synaptic pruning^18,45^ remain unknown. The human brain could conceivably adapt to distinct maturational tempos, and with only transient disparities, developmental trajectories might still converge within the normative range. While the distinctions in hierarchical brain organisation uncovered in our study may be transient, even these potentially short-term developmental discrepancies are essential to discuss since they could temporarily place the child outside the typical range and cause heightened stress levels. Subsequent investigations must undoubtedly address the longitudinal trajectories of brain development within cohorts characterised by diverse rates of maturation. It needs to be considered, however, that any such study faces a serious trade-off: observing long-term alterations due to accelerated puberty onset is more likely with age, but direct assessment of pubertal timing becomes less reliable with age. Although a long-term follow-up between puberty onset and adulthood seems necessary to complement our work, in the current study we have chosen to work with a cross-sectional sample to get a first glimpse on the impact of different maturational tempos on brain development.

With respect to the broader relevance of our findings, although the participants enrolled in the present study were deliberately chosen to fall within a normative range of maturational speed and were further screened for favourable socioeconomic circumstances, as well as the absence of developmental or neurological disorders, the findings bear more general implications, especially for clinical domains. We found an advantage of on-time maturation with respect to hierarchical brain organisation within our non-clinical cohort, indicating that significant deviations towards either accelerated or delayed maturational speeds might amplify disparities, leading developmental trajectories into the clinical spectrum. The acceleration of pubertal development is a growing concern today, particularly given the surge in medical referrals for young girls experiencing precocious puberty globally, with some instances manifesting as early as 6 or 7 years old^84–95^. Premature puberty carries the risk of compromised adult stature, psychosocial challenges, and potential health complications later in life^44,96,97^. The sudden peak in the lower left segment of the puberty onset age distribution curve (akin to that depicted in *Fig. 2a*), corresponding to instances of early puberty onset, suggests that there may be significant implications for the broader population involving a rapid secular trend. The possibility that developmental plasticity may not fully compensate for these changes is highly relevant. Secular trends in pubertal onset age persist across both developing and modern societies. While advancements in living conditions primarily account for trends in developing countries today^13–15,98,99^, negative anthropogenic influences, such as endocrine disrupting chemicals, have shaped the trend in developed nations in the last decades^98^. The recent pandemic has emerged as a novel potential factor influencing the age of pubertal onset, and it is essential to further investigate its impact on the general population.

In present times, within the most affluent nations, important developmental events, such as birth and puberty can be artificially timed, inducing intraspecies heterochrony which denotes variations in the timing of developmental events. Genetic-induced heterochrony introduces natural variability throughout evolution^100–102^. It remains uncertain whether anthropogenic-induced heterochrony (e.g., accelerated trends in pubertal timing, or artificially delayed puberty) will yield compromised mental functioning and health, or foster adaptive diversity in successive generations. Therefore, our study seems to be very timely, and can be considered as a first step towards investigating the potential outcomes of “anthropogenic heterochrony”. Mental health problems have surged following successive lockdowns, demonstrating the psychological toll on adolescents worldwide^57–60^. The findings of our study will aid in devising interventions more attuned to developmental contexts, thus lowering the risk of developing disabling psychopathologies in future generations.

## Methods

### Participants

The study is limited to investigating adolescent and emerging adult females, a deliberate choice made to disentangle the impacts of maturation and chronological age on brain development. Inclusion of both genders would have hindered in-depth analysis within the narrow temporal windows of biological and chronological age depicted in *Fig. 2b*. Given the delayed male pubertal onset^32,78^, a significantly larger sample would be required. We opted for inclusion of females since menarche age provides valuable supplementary data affirming the accuracy of bone age assessments in adolescents.

Participants were recruited via school contacts and online advertisements from Budapest, Hungary. Parents and/or participants provided demographic, educational, and medical history details. One participant with attention deficit disorder was excluded.

In order to determine the optimal sample size, we performed power calculations for two-sample z-test for means. A sample size of N ≥ 25 was found to be required for a power of at least 0.8 to detect a 0.8 effect at p<0.05. We aimed for a sample size N=30 in the adolescent and emerging adult chronological age groups. This yielded 20 subjects in the adolescent maturity groups with a power of at least 0.7. Ultrasonic bone age screening was conducted, and adolescent participants were categorised into BA/CA-defined bins, ensuring non-overlapping maturity groups: average (CA-0.5≤BA≤CA+0.5), accelerated (CA+1.5≥BA>CA+0.5), and decelerated (CA-1.5≤BA<CA-0.5). A pilot study revealed that around 20% of adolescents exhibited accelerated or decelerated maturation. Thus, 200 participants were invited (100 per CA group) for BA screening, aiming for ≥10 participants per bin. We excluded extreme cases (BA>CA+1.5, BA<CA-1.5) to avoid endocrinological complexities. This initial process resulted in a total of 65 participants between 12 and 14 years of age. However, during EEG preprocessing (see *Procedure*) 9 recordings were excluded from the study due to excessive noise. To compensate for this, we conducted further, targeted bone age screenings, and recorded EEG data of participants who fit into the aforementioned bin criteria, aiming to maintain a nearly equal distribution of participants across bins. From this additional recruitment, five new EEG recordings were successfully added. Thus, 61 girls aged 12 to 14 were included in the final cohort as depicted in *Fig. 2b*. The cohort of emerging adult participants consisted of 30 subjects. EEG data of 26 participants was included after the preprocessing procedure, as 4 recordings were removed due to noise.

The percentage of postmenarche participants in both age groups was lowest in the decelerated groups (see the Open Science Framework link under *Data availability*), suggesting that bone age is indeed a reliable gauge of maturity in adolescence. Mean menarche age of adolescent participants was 12.06y (SD=0.72). 27 of 61 participants did not provide data on menarche as they have not reached that stage yet. 88.52% of mothers and 83.61% of fathers of adolescent participants had a university degree demonstrating high socioeconomic status among the participating families. The EEG data collection was part of a large-scale study, within which each adolescent participant was also administered the Wechsler Intelligence Scale for Children, 4th edition (WISC-IV)^34^, carried out a fine motor task^36^, and a Stroop test^35^.

The Hungarian United Ethical Review Committee for Research in Psychology (EPKEB) approved the study (reference number 2017/84). Written informed consent was obtained from all subjects and their parents. Participants were given gift cards (approx. EUR 15 value each) for their attendance. All research described in this paper was performed in accordance with the Declaration of Helsinki.

### Procedure

#### Bone age assessment

Body measurements were taken according to the protocol laid down in the International Biological Programme^103^ using standard instruments. Skeletal maturity assessment began with an anthropometric procedure that includes the measurement of full body and sitting height (GPM Anthropometer, DKSH, Switzerland Ltd, Zurich, Switzerland), as well as body weight (Seca digital scale). Ultrasonic bone age estimation was conducted using the Sunlight BonAge System (Sunlight Medical Ltd, Tel Aviv, Israel), scanning the speed of sound (SOS) between the distal radius and ulna epiphysis^104^. This bone structure reflects the level of maturation as well as it changes significantly during stages of physical growth. The BonAge device calculates bone age in years and months using software that accounts for ethnicity and gender. Using ultrasound for skeletal maturity assessment is safe for the subjects as opposed to ionising radiation-based techniques, while results obtained from these two types of procedure are highly correlated^40^.

Bone age assessment was performed on the left hand and wrist of participants in each case. Participants were instructed to position their hands between the transducers on the unit’s armrest. The device then attached to the wrist, applying pressure of approximately 500g. Transducers transmitted 750 kHz frequency ultrasound through the wrist to measure SOS. The transducer on the ulnar side emits ultrasound while the other transducer acts as a receiver. Measurement protocol was repeated five times for accuracy, each measurement lasting about 20 seconds. Measurements were completed either at the high schools of participants, or at the Research Centre for Sport Physiology at the University of Physical Education, Budapest. The entire procedure took 5-10 minutes per person. Trained assistants conducted the measurements, and a biological anthropologist has overseen the measurement procedure and data analysis.

Bone age assessment results are considered valid for 3 months. Subjects were invited for the resting state electroencephalography (EEG) recording session within this timeframe. If a subject participated in laboratory testing 3 months after bone age assessment, the procedure was repeated.

#### Resting state EEG data recording

Continuous EEG data was recorded using the HydroCel GSN 130, a 128-channel high density EEG (HD-EEG) system (Electrical Geodesics, Inc., Canada), with the Net Station Acquisition software (version 5.4) in a windowless, air-conditioned room insulated from external noise and electromagnetic interference. Before recording, participants were asked to remove all electronic devices, e.g. phones and smartwatches from their pockets and bodies. Following a brief guided relaxation, we collected two five-minute (300 second) long resting state recordings at a 1,000 Hz frequency sampling rate. We aimed to keep impedances below 5 KΩ on all channels. EEG data recording took place at the Laboratory for Psychological Research, Pázmány Péter Catholic University, Budapest.

In the first 5-minute long recording segment, participants were instructed to keep their eyes open, blink minimally and to maintain posture to minimise artefacts generated by muscle movement^105^. During the second 5 minutes of recording, participants followed the same protocol, but were instructed to keep their eyes closed. The two conditions are separated since they exhibit different characteristics, namely a mismatch in topography and activity levels^106^. In the present study, we performed all statistical analyses on closed-eye recordings.

#### EEG data preprocessing

EEG preprocessing was performed in MATLAB (The MathWorks, Inc.; version: R2021a Update 4), using the EEGLAB toolbox^107^ (version 2021.1). First, we applied a high-pass filter at 1 Hz (transition bandwidth 1 Hz; passband edge 1 Hz; cutoff frequency -6 dB 0.5 Hz; zero-phase non-causal filter) to eliminate low frequency noise and direct-current offset, and a 40 Hz low-pass filter (transition bandwidth 10 Hz; passband edge 40 Hz; cutoff frequency -6 dB 45 Hz; zero-phase non-causal filter) to remove 50 Hz line noise and high frequency components, using EEGLAB’s basic finite impulse response (FIR; version 2.4) filter. Channel locations were then imported into EEGLAB, the channel layout map was supplied by Electrical Geodesics, Inc. Following filtering steps, noisy channels, determined by two independent raters via visual inspection, were excluded manually. As last steps, we removed noisy epochs from the recordings using the Clean Rawdata (version 2.4) plugin, then re-referenced data to average reference. The same preprocessing procedure was implemented for all recordings.

### Data Analysis

#### Leading Eigenvector Dynamics Analysis

After preprocessing, Leading Eigenvector Dynamics Analysis (LEiDA)^75^, adapted to EEG data by our group, was employed to analyse dynamic functional connectivity^69,73^ in the developing brain. The steps described below were performed on individual EEG recordings and were uniform in all cases.

First, preprocessed EEG recordings were filtered with a 6th order Butterworth bandpass filter for alpha band (8-12 Hz) without introducing a phase shift, then downsampled to 100 Hz (intervals of 10 ms). Subsequently, signals were detrended and demeaned, followed by a Hilbert-transformation to obtain the phase of all channels at all timepoints *t*. The nominal recording duration of 300 s yielded a temporal sequence of up to 30,000 phase vectors per subject. However, the EEGLAB toolbox typically excised small parts of the sequence, so that average sequence length was 28,500 vectors (SD=1,600). During preprocessing, we observed a minority of electrodes with interrupted EEG signal during some or all of the recording (“bad channels”, range 0 to 12; see *EEG data preprocessing*). To obtain full-size (128×128) coherence matrices at all timepoints and for every subject, we developed a two-step approach termed “patching”. In the first step, we formed group-level lists of “bad channels” by combining noisy channels from all subjects in a group, specifically, the 6 bins of adolescent participants (depicted in *Fig. 2e*) and the group of adult subjects. For *n* remaining “good channels” shared by all subjects of the group, instantaneous phase coherence *PC_xy,t_* was computed at every time point for each electrode pair as

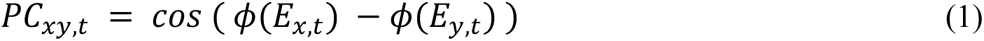

where Φ(*E_x,t_*) and Φ(*E_y,t_*) denote the instantaneous phase of electrode *x* and *y*, respectively, at each time point *t*. This resulted in a temporal sequence of *n* x *n* phase coherence matrices, sampled at 10 millisecond intervals, for each subject. The leading eigenvector (*1xn*) of each matrix was used to summarise the instantaneous pattern of phase differences between electrodes at every time point. Then, these vectors were clustered with Matlab’s built-in *k*-means clustering algorithm, assigning a cluster label to all timepoints in all subjects. In the second step, we patched the missing phase values of “bad channels” in all subjects. To this end, we replaced the missing phase values of a “bad channel” *x* at a time point *t* assigned to a particular cluster by randomly sampling a phase value from the ensemble of values recorded in other subjects of the same bin at the same electrode *x* and at time points assigned to the same cluster. Performing this procedure for all groups, subjects and “bad channels” permitted us to reconstruct full-size phase vectors (1×128) and full-size phase difference matrices (128×128). Full-size leading eigenvectors (1×128) were obtained from full-size phase difference matrices. Note that eigenvectors had the same dimension as the original phase vectors, but represented relative rather than absolute phase.

To identify patterns of phase differences that were common to all subjects – also referred to here as “brain states” and depicted in *Fig. 6* –, the leading eigenvectors at all timepoints and from all subjects were pooled and clustered via a modified *k*-means algorithm^108^ such as to obtain polarity invariant topographical maps. Through the modified *k*-means clustering algorithm, each vector is assigned to one of *k* clusters (or brain states) and is labelled accordingly (see *Fig. 6*), allowing us to represent the continuous EEG activity as a trajectory of, and transition between, brain states. LEiDA was carried out in Matlab (version: 2021a). While the results presented here are based on 12 clusters, analyses were also conducted using 6, 8, and 10 clusters and 12 clusters specific to each unique group (see the Open Science Framework link under *Data availability*). The results of these additional analyses were consistent with those presented in the *Results* section. We chose to present the results based on 12 brain states since it provides the most detailed description of the data.

**Fig. 6.**
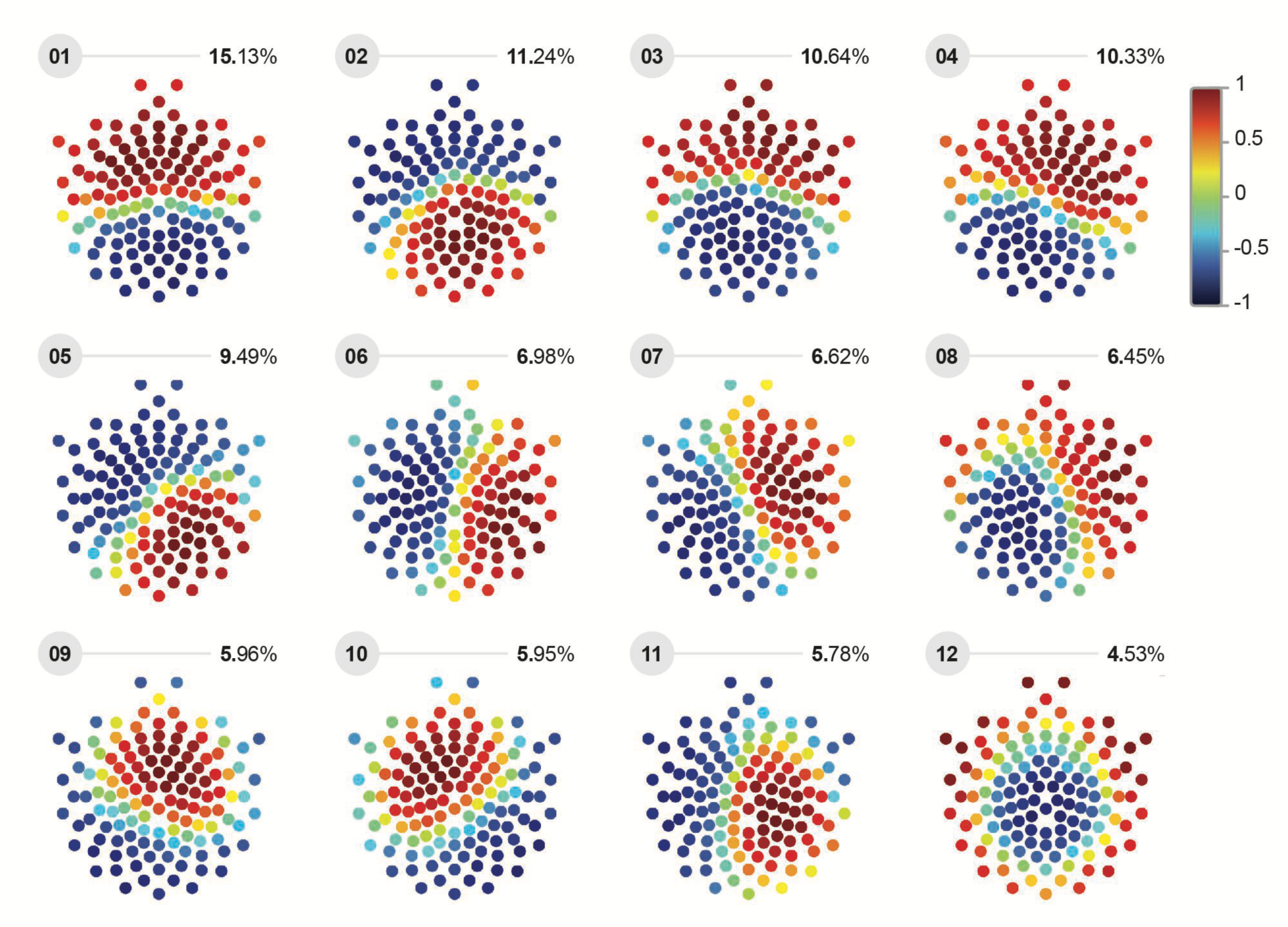
Brain state topographies. After initial preprocessing, EEG signals were analysed via Leading Eigenvector Dynamics Analysis (LEiDA). First, Hilbert transform was applied to each recording to acquire the phase of each electrode. Then, the instantaneous phase coherence between all electrode pairs was computed, yielding a 128×128 phase distance matrix for each time point *t*. Leading eigenvectors were extracted from the matrices via eigenvector decomposition. Following LEiDA, leading eigenvectors of all subjects were pooled, then clustered via modified *k*-means clustering to obtain topographically distinct connectivity patterns, which we refer to here as “brain states”. As leading eigenvectors from all subjects are combined, the twelve activity patterns are rendered uniform in each group. Brain states are ranked according to their dominance duration, expressed in percentages accompanying rank labels, with brain state number 01 accounting for the largest percentage of total activity, and brain state number 12 the smallest. The colour scale denotes cosine similarity, with similarly coloured dots indicating electrodes with congruent signal phases. Accordingly, values of 1 (dark red) and -1 (deep blue) represent similar and dissimilar (opposite) phases, respectively. Note that electrodes with similar and dissimilar phases form two spatially contiguous subsets.

A representative example for a sequence of successive patterns of phase differences (brain states) is shown in *Fig. 7a.* The average duration of brain states was 125 ms, with a standard deviation of ±0.4 ms over groups (range 12 ms to 13 ms) and ±1.5 ms over subjects (range 100 ms to 170 ms). However, the durations of individual brain states were quite variable (range 50 ms to 500 ms), with the average standard deviation over subjects being ±9 ms (range: 7 ms to 14 ms).

**Fig. 7.**
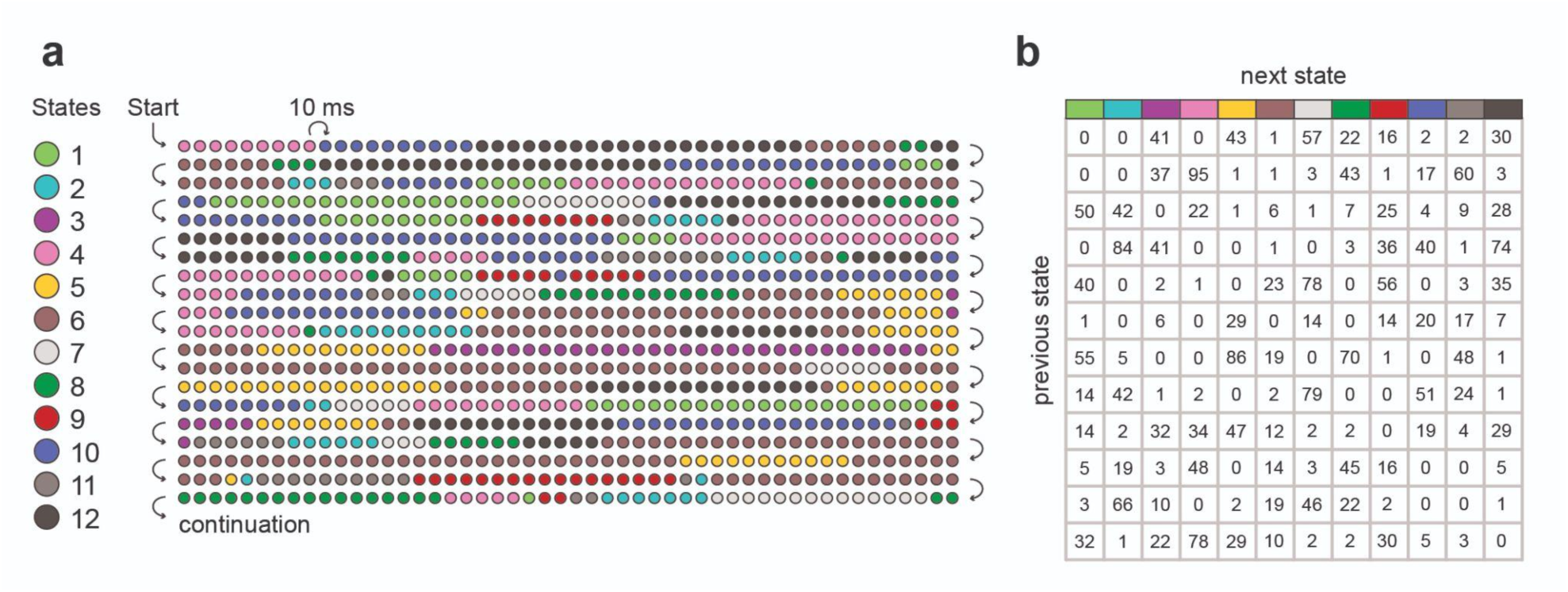
Transitions between brain states. **a,** Brain states numbered 1 to 12 are represented here by coloured disks. Only the initial 10s of a recording lasting 284 s are shown, specifically, the initial 1,000 leading eigenvectors (observed at 10 ms intervals) in a sequence of 28,400 eigenvectors. In this particular example, the average duration of brain states was 98 ms (i.e., 9.8 consecutive eigenvectors were assigned to the same state, on average). **b,** Number of observed transitions between different states in the entire sequence of 28,400 states. Previous states are represented by rows, next states by columns. Table entries indicate how often each particular transition was observed. Entropy production quantifies the degree of asymmetry in this matrix.

#### Entropy production calculation

To quantify asymmetry in the temporal dynamic of state space trajectories, we transformed the temporal sequence of brain states (*Fig. 7a*) of each subject into a table of transition counts *N_ij_* by removing diagonal entries (remaining in the same brain state) and by counting the observed number of forward and reverse transitions between pairs of different brain states (i,j). A representative example table is shown in *Fig. 7b*. To avoid singularities in the subsequent computation of entropy production, all counts were incremented by one, *N’_ij_=N_ij_+1*. Any effect of this change was nullified by the noise floor correction (see below). The corresponding transition probabilities were then obtained as *P_ij_=N’_ij_/Σ_ij_ N’_ij_*. To quantify the asymmetry in the pattern of state transitions, deemed indicative of the level of hierarchical organisation, we computed entropy production (also termed relative entropy^70^) for each subject as the Kullback-Leibler (KL) divergence:

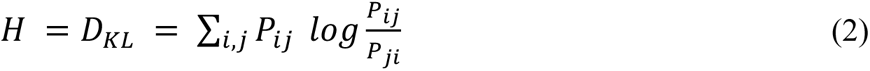

where *D_KL_* and *H* denote the total entropy production (i.e., the asymmetry between forward and reverse transition probabilities in brain state switching), *P_ij_* denotes the transition probability from brain state *i* to brain state *j*, while *P_ji_* expresses the probability of switching from brain state *j* to brain state *i*.

To estimate the variance of the observed entropy values, we generated 1,000 synthetic transition trajectories as Markov chains from the transition probability table observed for each subject via MATLAB’s two built-in functions, one of which creates discrete-time Markov chains and the other simulates Markov chain state walks. Synthetic trajectories had the same length as observed trajectories. Converting synthetic chains into entropy production values with *Eq. 2*, produced a distribution of a 1,000 entropy production values for each subject.

As the finite length of trajectories inflates entropy production values, we additionally performed a noise floor correction^70^. To this end, we synthesised a further 1,000 synthetic trajectories of the same length from a symmetric transition table (obtained by averaging the observed table with its transpose). Although perfectly symmetric transition probabilities should not produce any entropy, the resulting value distribution was positive, revealing inflation due to finite trajectory length. To obtain corrected entropy production values, we subtracted the mean symmetric-table entropy from the all asymmetric-table entropy values. From the distribution of corrected values, we obtained a mean and estimated variance of entropy production for every subject. Note that our procedure kept intact the temporal order of trajectories, taking subject-specific brain dynamics into account, improving on the simpler approach of Lynn and colleagues^70^. All calculations were performed in Matlab (version: 2021a).

#### Calculating group averages and confidence intervals

Group averages and variances of entropy production were computed by inverse-variance weighting. This was done to address the subject heterogeneity and to attenuate the influence of subjects with high variance on the weighted arithmetic mean. The following formulas were used:

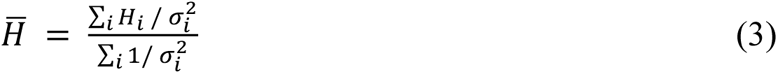

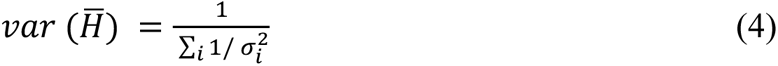

where 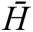 is the inverse variance weighted group average entropy production, *H_i_* denotes the mean observed entropy production for subject *i,* σ^i^_2_ denotes the observed variance for subject *i*, and 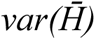 denotes the inverse variance weighted group variance. 95% confidence intervals for group-level mean entropy production was also computed based on 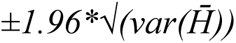. Group averages and confidence intervals were calculated in Matlab (version: 2021a).

#### Statistics

Statistical assessment of group differences was carried out using two-sample z-tests^109^, using the following formula:

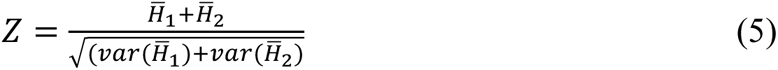

The significance level (*p*) in the two-sample z-test is determined as in the following:

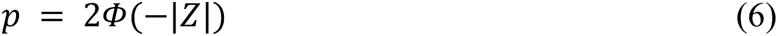

where Φ is the standard normal cumulative distribution function.

Results obtained from two-sample z-tests were reported with Hedges’ g effect size estimations. All statistical tests were carried out in Microsoft Excel Professional (version 2021).

### Data availability

Demographic and anthropological datasets, and EEG analyses are available at the Open Science Framework (OSF) platform at this address: https://osf.io/eqmwd/?view_only=d82d42bcb7a1481ca3ad33e721179523

## Acknowledgements

The authors thank T. Jáger for generating the figures. K. Utczás and Z. Tróznai for bone age data collection and analysis, G. Oláh for organising the participant cohort, L. J. Fehér for additional EEG data collection. We also thank the generous and amazing parents, adolescents and schools who participated in this project.

The project was supported by the National Research, Development and Innovation Office of Hungary (Grant K-134370 to I.K.) and by the Hungarian Research Network (HUN-REN-ELTE-PPKE Adolescent Development Research Group). The funders had no role in study design, data collection and analysis, decision to publish or preparation of the manuscript.

## Author Contributions

I.K. and F.G. conceptualised and designed the experiments; S.H. carried out data collection; M.L.K. and G.D. contributed theoretical analysis tools; M.C.M., J.B. and F.G. contributed analytical software; S.H., M.C.M., G.B., and J.B. carried out data analysis; F.G. carried out the statistical analysis; S.H. and I.K. drafted the manuscript. All authors discussed the results, contributed to the text, and approved the final version of the manuscript for submission.

## Table of Contents (ToC)

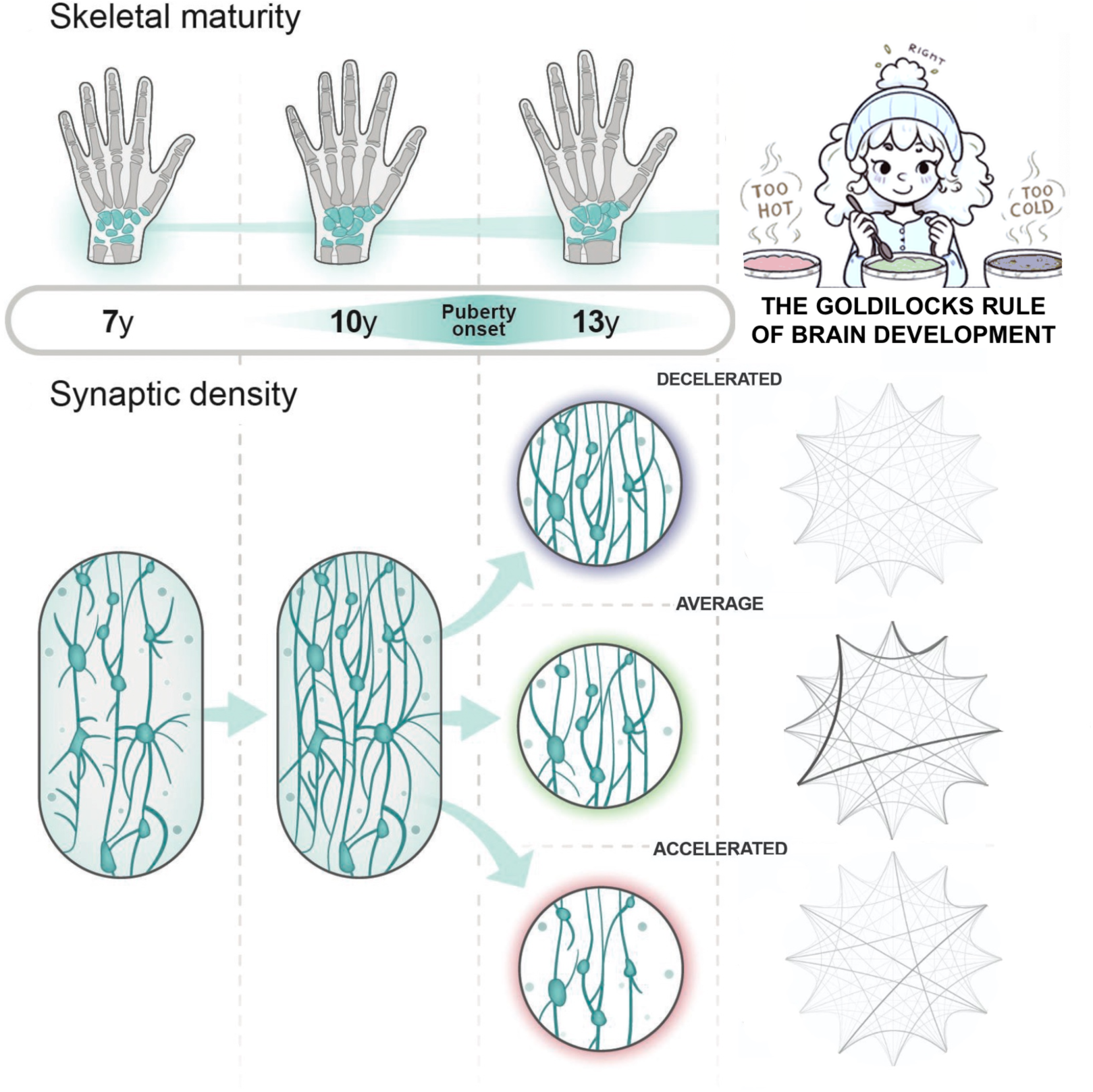

To reveal potential relationships between pubertal pace and the advancement of brain organisation, we analyse the connection between skeletal age-based maturation stages and hierarchical organisation in the temporal dynamics of resting-state EEG recordings (alpha frequency range). By adopting skeletal maturity as a proxy for pubertal progress and employing entropy production to measure hierarchical brain organisation, our findings indicate that an average maturational trajectory optimally aligns with cerebral hierarchical order.

